# hpGRISZ: a high-performance fluorescent biosensor for in vivo imaging of synaptic zinc dynamics

**DOI:** 10.1101/2025.10.06.680726

**Authors:** Hao Zhang, Manoj Kumar, Raquel Correia, Shengjie Yang, Wenyuan Huang, Carina Soares-Cunha, Patrick Andrew Cody, Yiyu Zhang, Xiaodong Tian, Shengyu Zhao, Mikhail Drobizhev, Ana João Rodrigues, Thanos Tzounopoulos, Hui-wang Ai

## Abstract

Despite the crucial role of synaptic Zn^2+^ in neurotransmission and neural processing, direct in vivo measurement of Zn^2+^ transients has remained challenging due to the limited responsiveness of existing fluorescent indicators. Here we report hpGRISZ (high-performance green indicator for synaptic Zn^2+^), an ultraresponsive turn-on green fluorescent biosensor engineered through iterative linker optimization, mutagenesis, and directed evolution. hpGRISZ exhibits exceptional brightness, thermostability, and a large fluorescence response (F/F_0_ ≈ 25) with micromolar affinity suitable for detecting extracellular synaptic Zn^2+^ release. We comprehensively characterized hpGRISZ in vitro, in mammalian cells, and in vivo. A membrane-anchored version of hpGRISZ traffics robustly to the cell surface under physiological conditions, where it retains strong responsiveness and supports wide-field, two-photon, and fiber photometry recordings in awake mice. These assays revealed synaptic Zn^2+^ dynamics across distinct brain regions and neuronal circuits. Together, our findings establish hpGRISZ as a powerful tool for dissecting zinc signaling in neural circuits and as a prototype for next-generation genetically encoded biosensors for in vivo imaging.

## Introduction

Zinc is the second most abundant transition metal in the human body and binds to ~10% of the proteome, serving essential catalytic and structural roles^1^. The levels of zinc ions (Zn^2+^) are compartmentalized and tightly regulated across organelles, vesicles, and extracellular spaces by a network of 24 transporters that maintain physiological Zn^2+^ distribution and balance^2,3^.

Disruption of Zn^2+^ homeostasis is linked to diverse pathologies, including immune and metabolic disorders^4–6^, as well as neurological diseases such as neurodegeneration,^3,7^ autism spectrum disorder^8^, depression^9^, epilepsy^10^, and Alzheimer’s disease^11^.

Beyond its structural and enzymatic roles, labile Zn^2+^—the pool of unbound, readily exchangeable zinc—acts as a neuromodulator^3^. Certain neurons load Zn^2+^ into synaptic vesicles and release it into the synaptic cleft, where it modulates pre- and post-synaptic receptors and ion channels in a manner dependent on Zn^2+^ concentration, timing, and spatial distribution^12–18^.

Despite its established role in synaptic signaling, knowledge of Zn^2+^ release and action has relied heavily on cell culture models, ex vivo preparations, or indirect inference from Zn^2+^ chelators—approaches that may fail to capture true physiological dynamics^19^. Direct measurement of Zn^2+^ signals in the intact brain has been limited by the scarcity of suitable tools, as existing indicators often lack the sensitivity, dynamic range, stability, appropriate affinity, or genetic targetability needed for in vivo imaging^19,20^. There is a critical need for advanced probes capable of capturing localized Zn^2+^ transients in behaving animals with high spatial and temporal fidelity^19,20^.

Synthetic and chemogenetic fluorescent Zn^2+^ indicators are valuable tools for studying zinc biology^19,21,22^, but genetically encoded fluorescent Zn^2+^ indicators (GEZIs) offer unique advantages for in vivo neuroscience, including cell-type-specific genetic targeting, stable expression without dye loading, and facile dissemination as DNA constructs^23^. Förster resonance energy transfer (FRET)-based GEZIs have provided a range of affinities and subcellular targeting options but are relatively bulky, spectrally broad, and typically display limited dynamic ranges^19,24-28^. Single-fluorescent protein (FP) GEZIs improved dynamic range and spectral simplicity^29–34^, yet they still exhibit limited responsiveness, suboptimal photophysical properties, mismatched affinities, and/or poor membrane targeting at physiological conditions (**Table 1**). Among green GEZIs, GZnP3 is the most responsive turn-on sensor but shows only a modest response at the mammalian cell surface^29^; designed for cytosolic Zn^2+^ detection, its affinity is also too high for reporting synaptic Zn^2+^ release^3^. Our previously reported ZIBG2 displayed low responsiveness and poor thermostability, precluding in vivo use^30^. To date, the only GEZI successfully applied in mice is FRISZ, a far-red sensor we developed earlier^33^; however, it has a modest dynamic range, is prone to aggregation in vivo, and has exhibited unreliable performance under two-photon imaging. These limitations underscore the need for a bright, thermostable, turn-on sensor optimized for reliable in vivo performance across diverse recording modalities.

**Table 1.** Performance comparison of hpGRISZ with representative genetically encoded Zn^2+^ indicators.

| Indicator | Type | Ex/Em<sup>a</sup><br>(nm) | Molecular<br>Brightness<sup>b</sup><br>(mM<sup>-1</sup>·cm<sup>-1</sup>) | K<sub>d</sub><sup>c</sup><br>(μM) | Dynamic<br>range<sup>d</sup><br>(fold) | T<sub>m</sub><sup>e</sup><br>(°C) | In vivo imaging? |
| --- | --- | --- | --- | --- | --- | --- | --- |
| **Green fluorescent Zn<sup>2+</sup> indicators** |  |  |  |  |  |  |  |
| hpGRISZ | single-FP<br>turn-on | 499/516 | 23.6 | 12 (protein);<br>4.4 (cell<br>surface) | 25 (protein);<br>22 (cell<br>surface) | 65.6 | Yes (awake mouse; wide-field, two-photon, fiber<br>photometry) |
| GZnP3 | single-FP<br>turn-on | 488/512 | 10.8 | 0.0013<br>(protein) | 17 (protein);<br>4 (cytosol)<br>2.8 (cell<br>surface) | ND<sup>f</sup> | No |
| ZIBG2 | single-FP<br>turn-on | 495/510 | 16 | 0.28 (protein) | 5 (protein);<br>1.8 (cell<br>surface) | 44.1 | No |
| **Far-red fluorescent Zn<sup>2+</sup> indicators** |  |  |  |  |  |  |  |
| FRISZ | single-FP<br>turn-on | 606/654 | 4.5 | 8.8 (protein);<br>7.3 (cell<br>surface) | 7.0 (protein);<br>2.8 (cell<br>surface) | ND<sup>f</sup> | Yes (awake mouse; wide-field) |
| **FRET-based ratiometric Zn<sup>2+</sup> indicators** |  |  |  |  |  |  |  |
| ZapCY /<br>eZinCh /<br>eCALWY<br>families | Dual-FP<br>FRET<br>ratiometric | Donor/acceptor-<br>dependent (e.g.,<br>CFP/YFP: Ex<br>~435; Em<br>~475/525) | NA<sup>g</sup><br>(ratiometric) | Variable (pM<br>to μM) | Typically < 2<br>(R<sub>max</sub>/R<sub>0</sub>) in<br>protein or cells | ND<sup>f</sup> | No |

<sup>a</sup> One-photon excitation/emission maxima for purified protein, as reported. When small spectral shifts occur with or without Zn<sup>2+</sup>, the values for the Zn<sup>2+</sup>-bound state are given.
<sup>b</sup> Molar extinction coefficient × quantum yield, determined using purified proteins for single-FP indicators. For FRET-based ratiometric sensors, brightness is not directly comparable.
<sup>c</sup> Apparent Zn<sup>2+</sup> affinities under specified conditions (purified protein or cell-surface display) at physiological pH, determined from intensity measurements.
<sup>d</sup> Fold change in fluorescence intensity (F/F<sub>0</sub>) or emission ratio (R<sub>max</sub>/R<sub>0</sub>) between Zn<sup>2+</sup>-bound and Zn<sup>2+</sup>-free states under indicated conditions.
<sup>e</sup> Thermostability temperature at which a 1-hour incubation retains 50% of its initial brightness.
<sup>f</sup> ND, not determined.
<sup>g</sup> NA, not applicable.

Here, we present hpGRISZ (high-performance green indicator for synaptic Zn^2+^), an ultraresponsive turn-on fluorescent indicator engineered for reliable in vivo imaging of extracellular synaptic zinc dynamics. hpGRISZ combines high brightness, a large dynamic range, robust thermostability, and efficient membrane targeting, thereby overcoming key limitations of previous GEZIs. We comprehensively characterized its photophysical and functional properties in vitro, in mammalian cells, and in vivo across multiple brain regions using wide-field imaging, two-photon microscopy, and fiber photometry. We show that hpGRISZ enables stable detection of Zn^2+^ transients in the auditory cortex (ACtx) and basolateral amygdala (BLA), two forebrain regions enriched in zincergic neurons via expression of the vesicular ZnT3 transporter^35,36^. Our data reveals that synaptic zinc release is dynamic and spatiotemporally-restricted to specific behaviorally-relevant events.

## Results

### Engineering hpGRISZ, a bright and ultraresponsive Zn^2+^ sensor

Through iterative rounds of rational linker optimization, structure-guided and random mutagenesis, and directed evolution, we developed hpGRISZ (**Fig. 1AB**)—a bright, thermostable green fluorescent Zn^2+^ indicator with a 25-fold turn-on response and affinity tuned for detecting extracellular neuronal Zn^2+^ release.

**Figure 1.**
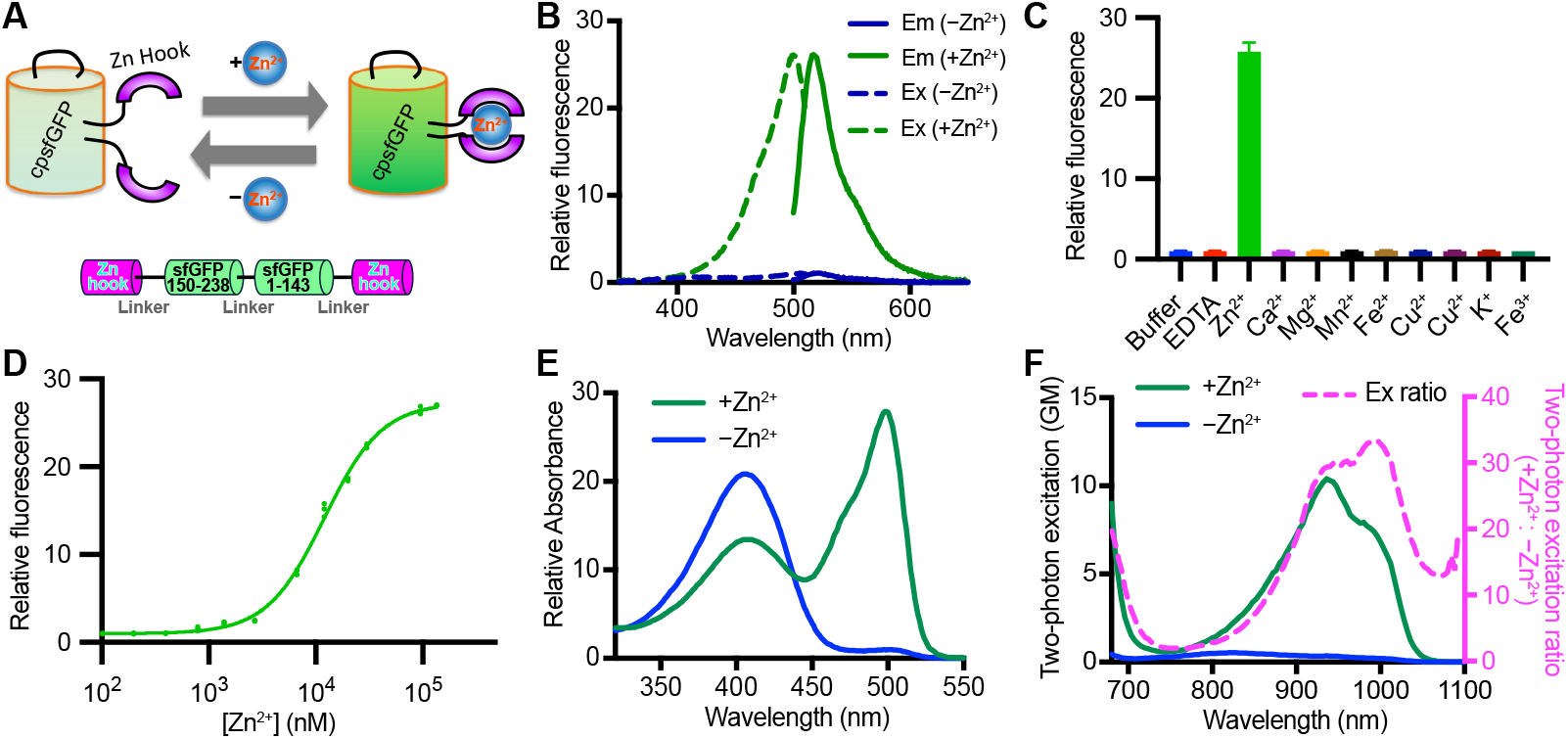
Mechanistic design and in vitro characterization of hpGRISZ. **(A)** Schematic of the Zn^2+^ sensing mechanism of hpGRISZ. Two Rad50 zinc hook motifs are fused to the N- and C-termini of circularly permuted superfolder GFP (cpsfGFP). Domain arrangement of the primary amino acid sequence is shown below. **(B)** Excitation (dashed lines) and emission (solid lines) spectra of hpGRISZ in the presence of 250 μM Zn^2+^ (green) or 250 μM EDTA (blue). **(C)** Fluorescence response of hpGRISZ to various metal ions (100 μM each). Data are presented as mean ± SD from three technical replicates. **(D)** Fluorescence response of hpGRISZ to increasing concentrations of Zn^2+^ (*K*_d_ = 12.0 ± 0.3 μM; n = 3 technical replicates). **(E)** One-photon absorbance spectra of hpGRISZ in the presence of 250 μM Zn^2+^or 250 μM EDTA. **(F)** Two-photon fluorescence excitation spectra of hpGRISZ in the presence of 250 μM Zn^2+^ (green) or EDTA (blue). The excitation ratio (Zn^2+^/EDTA, magenta) is plotted against excitation wavelength.

To promote robust folding and minimize redox sensitivity, we incorporated a cysteine-less, histidine-substituted variant of the *Pyrococcus furiosus* Rad50 zinc hook^37^ into a circularly permuted superfolder GFP (cpsfGFP) scaffold^38,39^ known for its folding stability (**Fig. S1**). Linker libraries of varying length and composition were cloned into a pTorPE periplasmic expression vector^33^ and expressed in *E. coli* at 37 °C. Colonies were imaged before and after spraying agar plates with Zn^2+^-chelating EDTA; hits showing fluorescence changes were advanced to liquid culture for plate-reader confirmation in cell lysates. This screening yielded GRISZ0.1, which, after multiple rounds of directed evolution, produced GRISZ0.3, a thermostable variant exhibiting ~2-fold (F/F_0_) turn-on to Zn^2+^, both as purified protein and when displayed on the surface of HEK293T cells.

We then modified the strategy by directly supplementing agar plates with Zn^2+^ to saturate the sensors and selected the brightest colonies for secondary screening in liquid culture. Screening a rationally designed, site-saturation mutagenesis library at four chromophore-proximal residues (147, 149, 256, and 263; **Fig. S2**), followed by iterative rounds of random mutagenesis, yielded variants with high apparent brightness and markedly enhanced dynamic range (**Fig. S3**). The top performer, GRISZ0.75, exhibited up to 50-fold dynamic range (F/F_0_) in vitro as a purified protein.

From GRISZ0.3 onward, promising mutants were tested using a pDisplay vector to anchor the sensor on the HEK293T cell surface. Although many mutants showed large dynamic ranges as purified proteins, responses were diminished on the mammalian cell surface; for example, GRISZ0.75 retained only a 2.2-fold (F/F_0_) response. Comparison with intermediate variants that performed better on the cell surface implicated residues 101 and 146 (**Fig. S2**) in modulating cell-surface performance. Accordingly, we constructed a combinatorial site-saturation library targeting both sites and identified GRISZ0.81 (G146K) and GRISZ0.82 (G146M), which showed improved cell-surface dynamic ranges (F/F_0_) of 6.4-fold and 4.3-fold, respectively.

We further hypothesized that discrepancies between bacterial and mammalian screening arose from the pTorPE plasmid, which encodes an N-terminal TorA signal peptide followed by a long linker. Inefficient TorA cleavage in *E. coli* may have left residual N-terminal residues on the expressed protein^40^. To test this, we appended these sequences to GRISZ0.75, GRISZ0.81, and GRISZ0.82 in their pDisplay constructs. The added segment improved performance across all three variants on the mammalian cell surface, with GRISZ0.82 exhibiting the strongest response enhancement (**Fig. S4**). We designated this N-terminally extended, high-performing construct as hpGRISZ **(Figs. S2 & S5)**.

### Exceptional photophysical performance in vitro

hpGRISZ was purified from *E. coli* using Twin-Strep affinity chromatography followed by size-exclusion chromatography, and exhibited a robust 25-fold fluorescence turn-on (F/F_0_) upon Zn^2+^ binding. The sensor exhibited single excitation and emission maxima; Zn^2+^ binding produced small blue shifts in the excitation maximum from 501 to 499 nm and in the emission maximum from 520 to 516 nm (**Figs. 1B** and **Table S1**). The fluorescence response was selective for Zn^2+^, with minimal changes upon addition of other common mono- and divalent metal ions under matched conditions (**Fig. 1C**). hpGRISZ exhibited a Zn^2+^-dependent fluorescence increase with an apparent dissociation constant (*K*_d_) of 12 μM (**Fig. 1D**), within the desired range for extracellular synaptic Zn^2+^ detection^3^. The sensor was also highly thermostable, retaining ~50% of its fluorescence after incubation at 65.6 °C for 1 hour (**Fig. S6**).

In the absence of Zn^2+^, hpGRISZ exhibited a dominant absorbance peak at ~406 nm with a minor shoulder near 500 nm (**Fig. 1E**). Upon Zn^2+^ binding, the short-wavelength band decreased while the long-wavelength band markedly increased, consistent with Zn^2+^-dependent chromophore deprotonation. Correspondingly, the extinction coefficient at ~500 nm increased from 1.8 to 25.6 mM−^1^·cm−^1^, and the quantum yield rose from 0.66 to 0.92 upon Zn^2+^ binding (**Table S1**). The Zn^2+^-free state showed a bi-exponential fluorescence lifetime decay [2.92 ns (83%) and 0.56 ns (17%)], whereas the Zn^2+^-bound state exhibited a mono-exponential decay (3.37 ns). Under two-photon excitation, hpGRISZ exhibited an excitation maximum at ~934 nm with a shoulder at ~996 nm; high two-photon brightness and Zn^2+^-dependent responses were observed across 880–1020, with a maximal ~33-fold enhancement (**Fig. 1F**).

### Robust membrane targeting and responsiveness in mammalian cells

hpGRISZ was introduced into a pDisplay plasmid containing an N-terminal Igκ leader sequence and a C-terminal PDGFRβ transmembrane (TM) domain. The resultant construct, which allows the expression of hpGRISZ on the mammalian cell surface, is referred as hpGRISZ-TM (**Fig. 2A**). When expressed at 37 °C on the surface of HEK293T cells, hpGRISZ-TM showed strong membrane localization and a robust Zn^2+^-dependent fluorescence increase. Membrane localization was assessed in HEK293T cells co-expressing cytosolic R-GECO^41^, a red Ca^2+^ sensor. hpGRISZ-TM exhibited clear membrane distribution distinct from the cytosolic R-GECO signal (**Fig. 2A**). Upon Zn^2+^ addition, hpGRISZ-TM fluorescence increased by 22-fold and was fully reversed by EDTA, a Zn^2+^ chelator (**Fig. 2B-D**). The response magnitude represented nearly an order-of-magnitude improvement over existing GEZIs (**Table 1**). R-GECO served as a negative control and showed no fluorescence change in response to Zn^2+^ or EDTA. Zn^2+^ titration on the cell surface revealed a concentration-dependent fluorescence response with an apparent *K*_d_ of 4.4 μM (**Fig. 2E**). Moreover, we assessed the response kinetics by simulating localized Zn^2+^ secretion with a 10-ms micropipette puff of Zn^2+^ solution (**Fig. S7**). The fluorescence half-times (*t*_0.5_) for rise (association) and decay (dissociation) were 191 ms and 422 ms, respectively, and the dissociation rate constant (*k*_off_) was 1.641 s^-1^— all within the range reported previously for FRISZ^33^.

**Figure 2.**
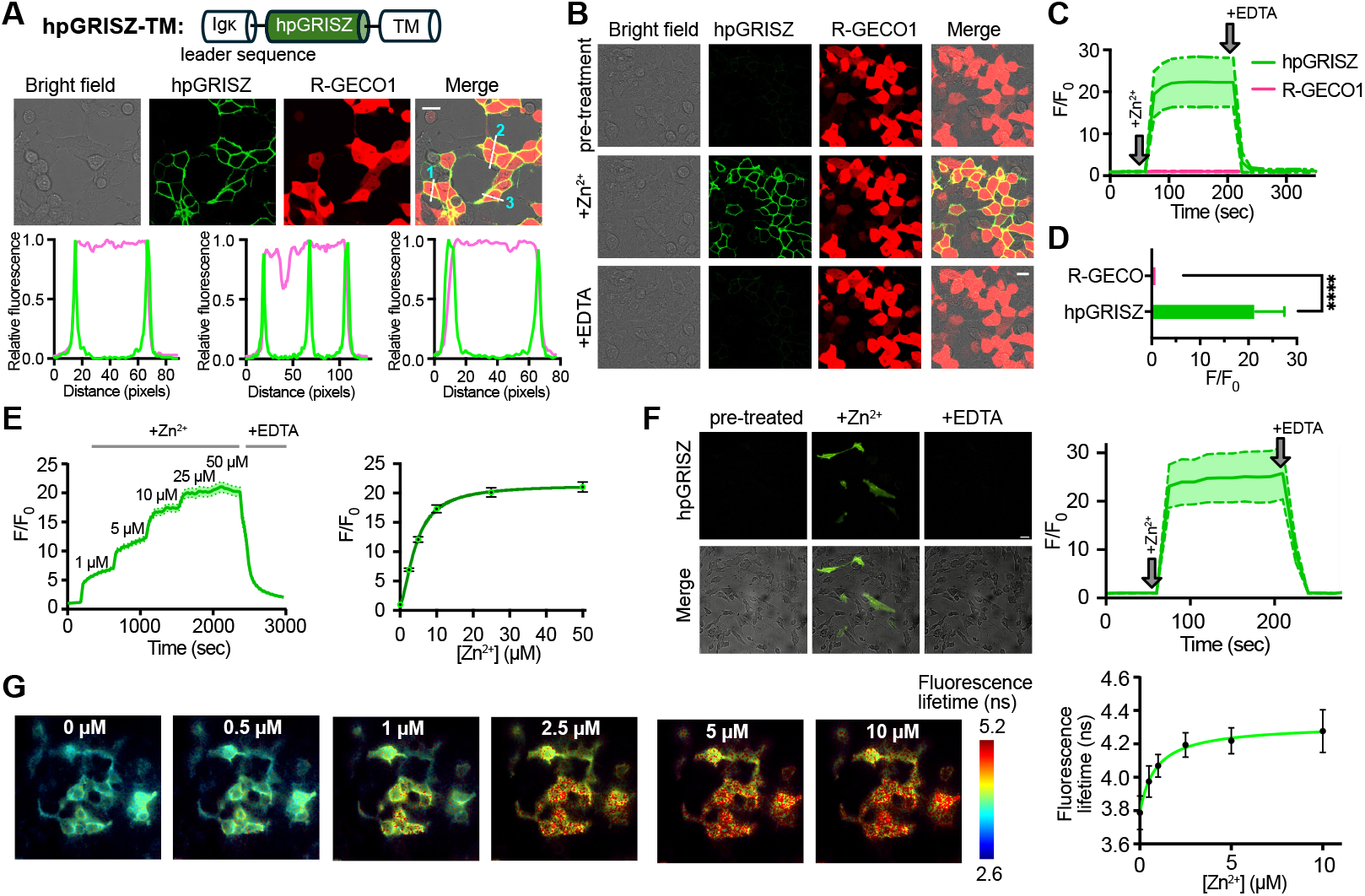
Characterization of hpGRISZ expressed at the mammalian cell surface. **(A)** Targeting of hpGRISZ to the surface of HEK293T cells. *Top:* Schematic of hpGRISZ-TM construct, including murine Igκ signal peptide and PDGFRβ transmembrane (TM) domain. *Middle:* Representative brightfield and fluorescence images of HEK293T cells co-transfected with hpGRISZ-TM and R-GECO1 (used here as a Zn^2+^-unresponsive cytosolic marker). Scale bar: 10 μm. *Bottom:* Fluorescence intensity profiles of hpGRISZ-TM (green) and R-GECO1 (magenta) along the indicated lines in the middle panel. **(B–D)** Time-lapse imaging of cells exposed to Zn^2+^ (100 μM) and EDTA (250 μM). *B:* Representative fluorescence images. Scale bar: 10 μm. *C:* Normalized fluorescence intensity traces over time. *D:* Quantification of maximal normalized fluorescence change (mean ± SD; \*\*\*\**P* < 0.0001, two-tailed unpaired t-test). **(E)** Dose-dependent fluorescence response of hpGRISZ at the HEK293T cell surface (*K*_d_ = 4.4 ± 0.3 μM; mean ± SEM from 77 cells). **(F)** Time-lapse fluorescence images (left) and intensity trace (right) of hpGRISZ expressed at the SH-SY5Y cell surface in response to Zn^2+^ and EDTA (mean ± SEM from 27 cells). Scale bar: 10 μm. **(G)** Dose-dependent fluorescence lifetime response of hpGRISZ at the HEK293T cell surface. *Left*: Pseudo-color lifetime images. *Right:* Corresponding dose-response curve (*K*_d_ = 0.82 ± 0.13 μM; mean ± SEM from 34 cells). Scale bar: 10 μm.

We next tested hpGRISZ-TM in the neuroblastoma SH-SY5Y cell line, where it localized correctly to the plasma membrane and retained a >20-fold fluorescence response to Zn^2+^ (**Fig. 2F**). Finally, we evaluated hpGRISZ-TM as a fluorescence lifetime sensor and found that it exhibited a Zn^2+^-dependent lifetime change of up to 0.49 ns, with an apparent *K*_d_ of 0.85 μM (**Fig. 2G**), supporting its potential suitability for quantitative Zn^2+^ measurements.

Together, our in vitro and cellular characterization establish hpGRISZ as a next-generation fluorescent Zn^2+^ indicator, combining high brightness, robust responses both in purified form and at the mammalian cell surface, appropriate Zn^2+^ affinity and response kinetics, exceptional thermostability, and compatibility with one- and two-photon excitation—representing a substantial advance over existing GEZIs (**Table 1**).

### Wide-field and two-photon imaging of hpGRISZ tracks sound-evoked ACtx Zn^2+^ dynamics in awake mice

Synaptic Zn^2+^ fine-tunes ACtx sound processing and supports conserved adaptation mechanisms, including deviant detection—where repeated stimuli evoke reduced neuronal responses while novel (deviant) stimuli trigger robust activity—and contrast gain control, which adjusts sensitivity to changing background sound statistics^17,18,42-45^. Thus, to evaluate hpGRISZ-TM as a tool for directly measuring synaptic zinc signaling in awake mice, we expressed hpGRISZ-TM in the ACtx to assess sound-evoked Zn^2+^ release (**Fig. 3A**). To achieve this, we generated a viral vector to express the sensor under the neuron-specific hSyn promoter (AAV2/9-Syn-hpGRISZ-TM). Viral injection into the ACtx resulted in robust expression, confirmed by wide-field epifluorescence imaging at 12–16 days post-injection (**Fig. S8A**). To assess sound-evoked Zn^2+^ responses, awake mice were presented with 100-ms broadband sound stimuli (6–64 kHz, 60–70 dB SPL) while fluorescence changes were recorded under a 4× objective. Spatial ΔF/F maps identified sound-responsive regions. Sound-evoked signals exhibited two components: a fast motion-related transient (< 0.3 s post-stimulus) and a slower (> 0.3 s) positive excursion (**Fig. S8A**). To determine the spatial origin of these components, we systematically swept 10×10 pixel ROIs at 5 pixel increments across the field of view and clustered responses by temporal similarity (**Fig. S8B**). This analysis revealed that fast negative excursions clustered along blood vessel edges (**Fig. S8CD**). Motion tracking of vessel displacement confirmed that these signals were artifacts, likely arising from bidirectional movements that resolved by ~ 0.3 s post-stimuli. When vessel motion obscured the underlying tissue, baseline fluorescence decreased (negative excursions), whereas displacement that uncovered previously hidden fluorescent tissue produced signal increases (positive excursions; **Fig. S8EF**). To attenuate these motion artifacts, which occurred within 300 ms of sound onset, we applied a low-pass Butterworth filter (see **Methods**). Using this analysis pipeline, we compared responses before and after local infusion of ZX1, a fast extracellular Zn^2+^ chelator^12^. ZX1 largely reduced the sound-evoked hpGRISZ-TM signals (**Fig. 3B**), whereas ACSF infusion had no effect (**Fig. 3C**), confirming that hpGRISZ-TM faithfully reports sound-evoked Zn^2+^ release. There were residual hpGRISZ responses after ZX1 infusion, consistent with previous findings suggesting incomplete elimination of Zn^2+^ by ZX1^17,33^.

**Figure 3.**
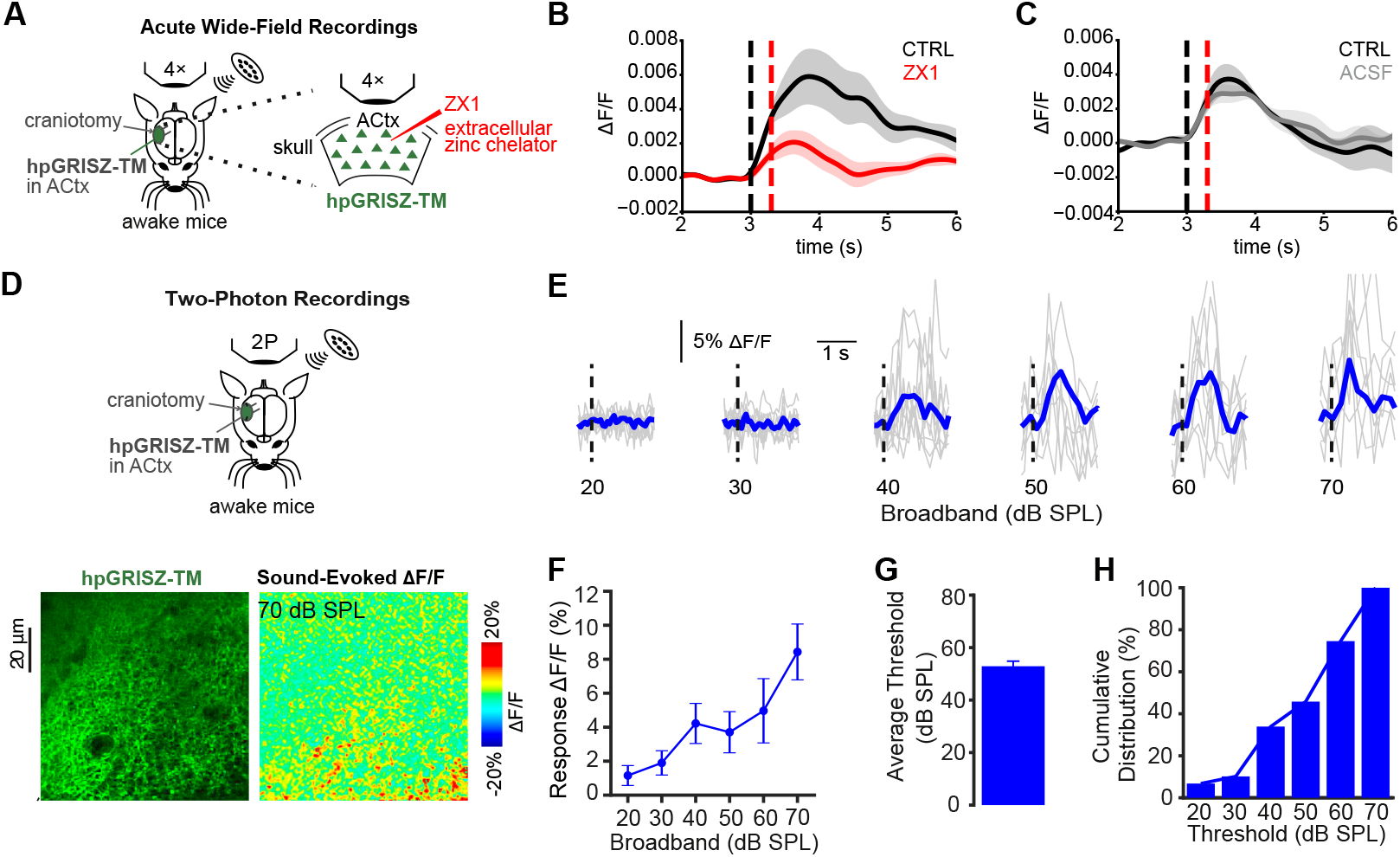
Wide-field and two-photon fluorescence imaging of sound-evoked Zn^2+^ signals using hpGRISZ-TM in awake mice. **(A)** Schematic of wide-field epifluorescence imaging setup targeting the auditory cortex (ACtx) in awake mice. **(B)** Average ΔF/F ± SEM before (CTRL, black) and after ZX1 infusion (red) following low-pass filtering (n = 7 animals). **(C)** Same as panel B, but before (CTRL; black) and after (gray) ACSF infusion (n = 4 animals). **(D)** Schematic of two-photon imaging setup for layer 2/3 neurons in ACtx. Representative fluorescence image shows hpGRISZ-TM expression, and sound-evoked response (ΔF/F) shows Zn^2+^ release hotspots. **(E)** Representative fluorescence traces of hpGRISZ-TM in response to broadband sound. Blue traces represent the average of individual trials (grey traces) **(F)** Average response amplitude of individual layer 2/3 neurons (59 neurons in 3 mice). **(G)** Average response threshold of individual layer 2/3 neurons. **(H)** Cumulative distribution of response thresholds.

To further image Zn^2+^ signaling at single-cell resolution, we performed two-photon imaging of layer 2/3 neurons in awake mice expressing hpGRISZ-TM. To monitor Zn^2+^-mediated fluorescence changes, broadband sound stimuli (6–64 kHz, 100 ms long) were delivered at intensities ranging from 20 to 70 dB SPL in 10-dB steps (**Fig. 3D, top**). We found scattered, sound-evoked green fluorescence increases, suggesting sparse synaptic Zn^2+^ release in cortical neurons (**Fig. 3D, bottom**). Automated analysis of the somatic regions of 335 neurons from three mice revealed that 59 cells with significant sound-evoked Zn^2+^ responses (**Fig. 3E**).

Responses were strongly sound-intensity-dependent, with an average threshold of 52.8±1.9 dB SPL and a broad threshold distribution (20–70 dB SPL) (**Fig. 3E-H**). Together, these results demonstrate that hpGRISZ-TM enables reliable in vivo detection of sound-evoked synaptic Zn^2+^ dynamics across mesoscale and single-cell levels, indicating that Zn^2+^ signaling in the ACtx is spatially sparse, cell-specific, and modulated by stimulus intensity.

### Fiber photometry recording of hpGRISZ reveals Zn^2+^ dynamics in the BLA of freely behaving mice

The vesicular zinc transporter ZnT3 is highly enriched in the BLA, a critical hub for aversive information processing^46^. Prior ex vivo studies demonstrated that synaptically released Zn^2+^ facilitates long-term potentiation at cortico-amygdala synapses, suggesting a role in persistent synaptic modifications underlying fear learning^35^. In line with this, ZnT3 knockout mice show normal fear conditioning with strong tone–shock pairings but exhibit reduced fear memory under weaker training^46,47^, implicating BLA Zn^2+^ signaling in modulating aversive responses.

To directly investigate Zn^2+^ signaling during aversive experiences, we used fiber photometry in freely moving mice to record hpGRISZ-TM signals in the BLA (**Figs. 4A**). A hybrid optofluidic cannula enabled simultaneous fluorescence recording and local infusion of either ZX1 or saline control solutions. Across three aversive paradigms, we observed robust, stimulus-locked Zn^2+^ transients. Looming sound^48^, which mimics an approaching predator, elicited clear increases in green fluorescence, which were abolished by ZX1 (**Fig. 4BC**). Individual trial heatmaps revealed consistent responses across animals, with area under the curve (AUC) significantly elevated during stimulus periods compared to baseline (**Fig. 4DE**). Similar Zn^2+^ transients were detected in response to a 20 kHz pure tone (**Fig. 4F-I**), an ultrasonic frequency previously shown to be aversive to rodents, in part because it overlaps with alarm vocalizations^49^. The strongest responses were evoked by foot shock (0.25 mA), producing large fluorescence increases that were substantially attenuated by ZX1 treatment (**Fig. 4JK**). Heatmap visualization confirmed the reliability and magnitude of shock-induced Zn^2+^ signals across animals, with AUC analysis again demonstrating significant ZX1 sensitivity (**Fig. 4LM**). Across all three paradigms, ZX1-treated mice displayed small negative fluorescence responses to stimuli (**Fig. 4CGK**), likely reflecting transient acidification associated with synaptic vesicle release. Notably, these pH-related artifacts produced signal changes in the opposite direction of Zn^2+^-evoked fluorescence increases, further supporting the robustness of hpGRISZ for detecting synaptic zinc dynamics.

**Figure 4.**
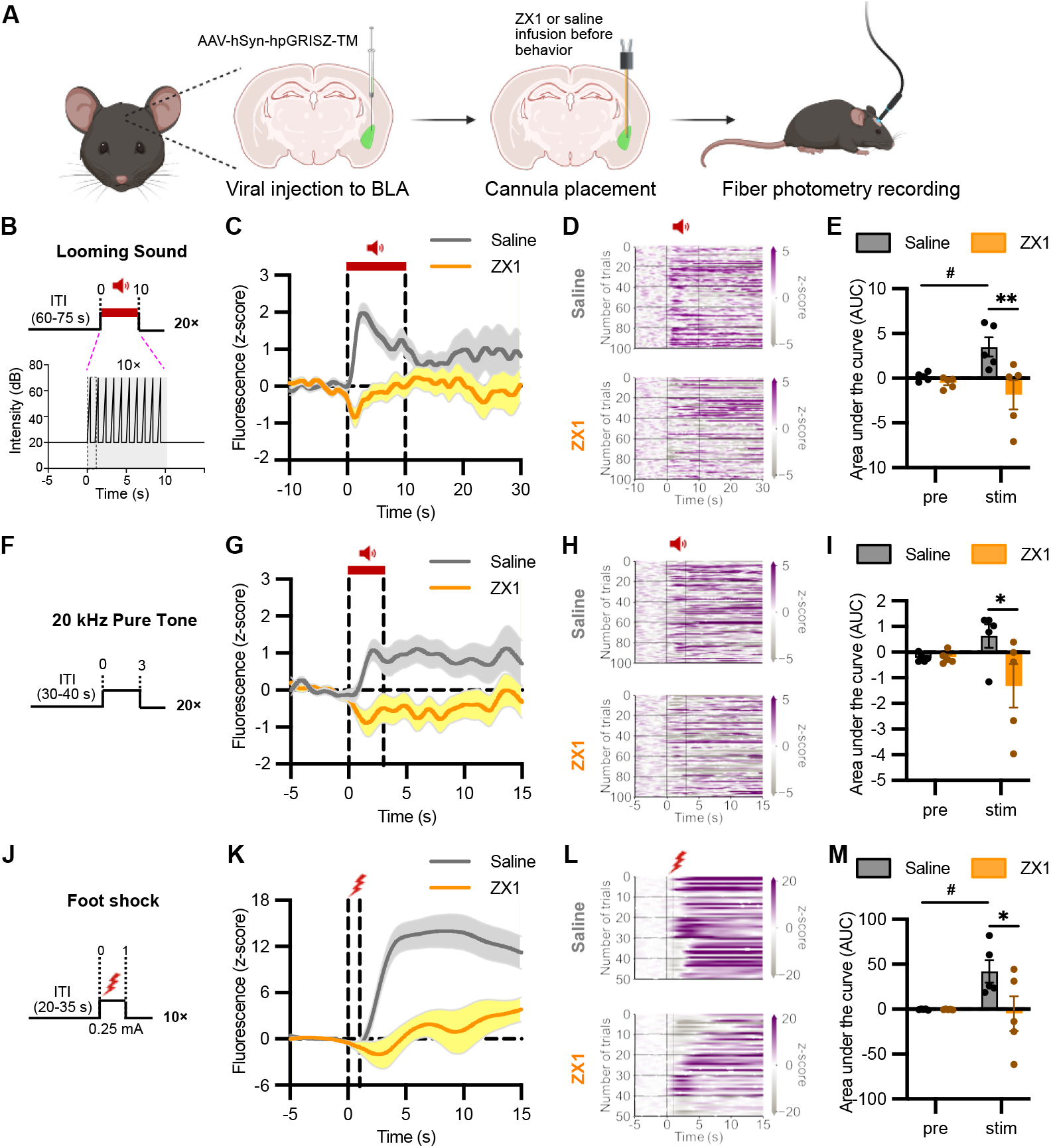
Fiber photometry recording of Zn^2+^ dynamics in the basolateral amygdala (BLA) in freely moving mice during aversive stimuli. **(A)** Experimental overview. Mice received BLA injections of hpGRISZ-TM virus and hybrid optofluidic cannula implantation for simultaneous compound infusion and fiber photometry. Immediately before each session, mice received intra-BLA injections of saline or ZX1 (200 μM). Schematic of looming sound protocol (n_saline_ = 5; n_ZX1_ = 5), consisting of 20 trials (10 s each) with a 0.4 s ramping sound after a 0.6 s delay. Inter-trial intervals (ITIs) were 60–75 s. **(C)** Looming sound evoked an increase in hpGRISZ-TM fluorescence in saline-injected mice, abolished by ZX1. **(D)** Heatmaps of z-scored fluorescence changes. Each horizontal line represents one trial; trials are grouped by animal, with horizontal black lines separating individual animals. Stimulus onset indicated by vertical black lines. Fluorescence signals were normalized to baseline on a trial-by-trial basis. **(E)** Quantification of area under the curve (AUC) during pre- and stimulus (stim) periods. **(F)** Schematic of 20 kHz pure tone protocol (n_saline_ = 5; n_ZX1_ = 5): 20 trials (3 s each) with ITIs of 30–40 s. **(G)** Fluorescence increased during tone exposure in saline-injected animals, but the increase was suppressed after ZX1 infusion. **(H)** Heatmaps of fluorescence across trials and animals during pure tone trials. **(I)** AUC analysis for pre- and stimulus periods during tone exposure. **(J)** Schematic of foot shock protocol (n_saline_ = 5; n_ZX1_ = 5): 20 mild shocks (0.25 mA), ITIs of 20–35 s. **(K)** Foot shock elicited a strong fluorescence increase in the BLA, reduced by ZX1. **(L)** Heatmaps of fluorescence across trials and animals during foot shock trials. **(M)** AUC analysis of pre- and stimulus periods during shock trials. Data are represented as mean ± SEM (* or # *P* ≤ 0.05; ** *P* ≤ 0.005; two-way ANOVA followed by Šídák’s multiple comparisons tests).

Together, these experiments establish that the BLA exhibits pronounced extracellular Zn^2+^ transients during diverse aversive stimuli. Their activation by stimulus presence and suppression by extracellular Zn^2+^ chelation support a synaptic origin, highlighting a critical role for Zn^2+^ signaling in amygdala circuits underlying aversive processing.

## Discussion

The development of hpGRISZ overcomes multiple limitations of prior GEZIs and enables robust, high-resolution imaging of extracellular synaptic Zn^2+^ dynamics in vivo. By combining thermostable scaffold selection, targeted linker and chromophore-proximal mutagenesis, and iterative bacterial/mammalian screening, we generated a sensor with high brightness, large dynamic range, thermostability, and suitable affinity for synaptic Zn^2+^ dynamics. Importantly, hpGRISZ maintains strong responsiveness when anchored to mammalian cell surface and supports stable wide-field, two-photon, and fiber photometry recordings in awake animals.

These advances extend the utility of prior sensors and provide the first demonstration of synaptic Zn^2+^ imaging beyond wide-field approaches, enabling single-cell resolution and multi-region recordings in behaving mice.

Several important considerations emerge from our sensor engineering efforts. First, the mutations that enhanced the sensor performance are distributed throughout the protein (**Fig. S5**), making it difficult to rationalize how each individual substitution contributes to the overall functional outcome. This underscores the complex and often non-intuitive relationship between sequence variation and performance in genetically encoded sensors. Second, there is a clear disconnect between sensor properties measured in purified proteins and those observed at the cell surface, and the determinants of cell-surface performance remain incompletely understood. While thermostability appears to play a role, it is unlikely to be the sole factor. For instance, GRISZ0.75, GRISZ0.81, and GRISZ0.82 differ by only a single residue (146, projecting outward from the sfGFP β-barrel), yet show markedly different behaviors both as purified proteins and in mammalian surface-expression contexts (**Fig. S4**). Similarly, GRISZ0.82 and hpGRISZ (GRISZ0.82 with additional N-terminal sequences) exhibit comparable fluorescence thermostability but diverge substantially in their cell-surface responses. It is worth noting that the AlphaFold3 structural model of hpGRISZ suggests potential interactions between these N-terminal sequences and other regions of the protein (**Fig. S5**). Together, these findings highlight the complexity of engineering reliable surface-targeted sensors and indicate that multiple, as yet undefined, factors contribute to functional performance in vivo. At present, directed evolution and empirical testing remain the most reliable approaches for developing effective sensors.

While wide-field imaging offers a valuable overview of neuronal activity across a large cortical area encompassing multiple layers, it lacks the resolution needed to reveal heterogeneous synaptic Zn^2+^ dynamics within individual neurons. Given zinc’s established role in shaping ACtxs sound processing and adaptation in a context-, cell-type- and layer-specific manner,^17,18,42^ resolving Zn^2+^ signaling with high precision is essential. hpGRISZ addresses this need by enabling single-cell resolution imaging of synaptic Zn^2+^ in a layer-specific manner, thereby providing a new window into how Zn^2+^ contributes to the fine-tuning of sound-evoked cortical activity. Looking forward, Cre-dependent AAV expression of hpGRISZ-TM will make it possible to selectively target excitatory versus inhibitory neurons, enabling detailed, cell type-specific analyses of sound-evoked Zn^2+^ dynamics.

An important consideration for in vivo Zn^2+^ imaging is distinguishing real signals from motion artifacts. We identified brief, sound-evoked motion artifacts with faster kinetics (<300 ms) than genuine Zn^2+^ responses, particularly near blood vessels where lower baseline fluorescence enhances these artifacts. While the precise mechanisms underlying these artifacts remains to be explored, neurovascular coupling or sound-evoked muscle contractions may contribute to these signals. Critically, the distinct temporal differences between motion artifacts and Zn^2+^ responses allowed reliable discrimination, enabling hpGRISZ-TM to accurately capture sound-evoked Zn^2+^ dynamics across both wide-field and two-photon modalities. This temporal discrimination represents a key methodological consideration for future in vivo imaging studies.

Our in vivo studies highlight a fundamental role for Zn^2+^ signaling across distinct brain regions. In the ACtx, Zn^2+^ responses were tuned by stimulus intensity, supporting a role in sound processing, while in the BLA, robust Zn^2+^ transients were consistently evoked by aversive stimuli, supporting a role in associative learning and fear-related processing. Together, these results provide evidence that synaptic zinc serves circuit-specific functions—modulating sensory computations in neocortical regions and regulating plasticity and aversive behaviors in limbic circuits. The robust and stimulus-dependent Zn^2+^ signals observed in these brain regions align with zinc’s known actions at glutamatergic and GABAergic receptors and its established role in regulating synaptic plasticity, including long-term potentiation (LTP) and long-term depression (LTD)^50–52^. Prior studies using constitutive ZnT3 knockout mice have implicated zincergic signaling in fear conditioning and extinction, but interpretations have been complicated due to potential compensatory mechanisms^46,47^. Future work combining hpGRISZ with the recently reported conditional ZnT3 knockout models^53^ could provide the causal resolution needed to define zinc’s role in circuit function and behavior—an avenue we are actively pursuing.

One long-standing challenge in zinc biology is the accurate determination of Zn^2+^ concentrations in intact biological systems^54,55^. Complex microenvironments, rapid signaling transients, and the coexistence of bound and labile Zn^2+^ pools hinder precise quantification. Our fluorescence lifetime measurements in mammalian cells provide proof-of-principle that hpGRISZ can enable quantitative assessment of Zn^2+^. Because fluorescence lifetime is probe-concentration-independent, this approach holds strong potential for standardized Zn^2+^ measurements in vivo. While our current results are preliminary and limited to cultured cells, they highlight a promising direction for applying hpGRISZ-TM in complex tissues in vivo, a critical step for further mechanistic insight.

In addition, the robust folding scaffold of hpGRISZ may support innovative engineering strategies that extend beyond conventional sensor design. One intriguing possibility is the development of split-sensor variants that reconstitute only at cell–cell interfaces, enabling synapse-targeted detection with sub-synaptic specificity^56^. Such next-generation tools would open the door to direct visualization of Zn^2+^ dynamics at defined synaptic contacts, providing unprecedented mechanistic insight into how Zn^2+^ contributes to circuit-level signaling.

Furthermore, the versatility of hpGRISZ across imaging platforms makes it well suited for integration with complementary methods such as optogenetics, electrophysiology, and behavioral paradigms. This performance profile also opens avenues for investigating zinc’s role in neurological and psychiatric disorders. Altered synaptic Zn^2+^ signaling has been implicated in in conditions such as Alzheimer’s disease, depression, epilepsy, schizophrenia, and traumatic brain injury, yet mechanistic links remain elusive^7^. hpGRISZ could be deployed in disease models to monitor region-specific zinc dysregulation in real time, potentially revealing early biomarkers of progression or treatment response. Beyond neuroscience, hpGRISZ may also be adapted to interrogate zinc biology in other tissues, such as pancreatic islets—where β-cell Zn^2+^ release is tied to insulin dynamics^6^—extending its utility to systemic zinc regulation.

## Methods

### Ethical statement

All experiments were conducted in accordance with relevant ethical standards. Animal procedures were approved either by the Institutional Animal Care and Use Committee (IACUC) at the University of Pittsburgh (Protocol # 23114004 and 23063309) or by the Animal Welfare and Ethics Committee (ORBEA) of the ICVS (Protocol # 009-2020) and the Portuguese authority DGAV (Protocol #8332-2021). C57BL/6J mice (The Jackson Laboratory, stock #000664) were housed under standard conditions, including a 12-hour light/dark cycle, ambient temperatures of 20–25 °C, and humidity maintained at 40–60%. Animals were randomly assigned to experimental groups to ensure inclusion of both sexes. Portions of the text were edited for clarity and grammar using ChatGPT (OpenAI), with all content reviewed and verified by the authors.

### Library construction and bacterial screening

Random mutagenesis was performed using error-prone PCR (EP-PCR) with Taq polymerase, MnCl_2_, MgCl_2_, and an unbalanced dNTP mix as previously described.^33^ Site-directed and site-saturation mutagenesis were carried out using mutagenic primers. Amplified fragments were cloned into a modified pTorPE vector (pTorPE-FRISZ^33^, Addgene #176887), which contains an N-terminal TorA periplasmic export sequence followed by a linker, and a C-terminal Twin-Strep-tag. Gene libraries were used to transform *E. coli* DH10B, which were plated on selective 2xYT agar containing ampicillin (100 μg/mL) and L-arabinose (0.02%, w/v). In the early screening rounds, fluorescence responses of bacterial colonies were examined by imaging before and after applying 2 mM EDTA (a Zn^2+^ chelator) using a fine mist sprayer. In later rounds, ZnCl_2_ (100 μM) was directly added to saturate the sensors and facilitate selection of the brightest colonies. Promising clones were then cultured in 96-well plates at 37 °C overnight (~18 h) and lysed. The clarified supernatants were assayed in 96-well format by adding Zn^2+^ (250 or 50 μM) or EDTA (100 μM). Fluorescence responses were measured on a BioTek Synergy Mx microplate reader, and variants with strong signals were advanced to subsequent rounds of engineering or detailed characterization.

### Protein expression and characterization

Single colonies of *E. coli* DH10B harboring the plasmids were grown in 500 mL 2xYT medium (100 μg/mL ampicillin, 0.02% L-arabinose) at 37 °C, 250 rpm for 24 h. Twin-Strep-tagged proteins were purified using Strep-Tactin Sepharose (IBA Lifesciences) followed by size-exclusion chromatography on a HiLoad 16/600 Superdex 200 pg column (Cytiva, ÄKTA system) and eluted in HEPES buffer (150 mM HEPES, 100 mM NaCl, pH 7.4). Fresh proteins were diluted in assay buffer for in vitro measurements on a BioTek Synergy Mx plate reader. For spectral scans, 100 nM protein with 250 μM Zn^2+^ or 250 μM EDTA was used; excitation spectra (350–500 nm) were collected with emission at 520 nm, and emission spectra (500–650 nm) with excitation at 480 nm. Endpoint measurements used excitation/emission at 495/515 nm. Metal selectivity was tested by adding 100 μM of various salts (ZnCl_2_, CaCl_2_, MgCl_2_, MnCl_2_, FeSO_4_, CuSO_4_, CuI, KCl, FeCl_3_) to 100 nM protein, and fluorescence was recorded immediately. Zn^2+^ titrations (101 nM–1.35 mM) were fitted to the Hill equation. For thermostability assays, 100 nM protein in 100 μM EDTA was incubated at various temperatures for 1 h in a thermal cycler, cooled to room temperature, and endpoint fluorescence was recorded.

### Photophysical and two-photon spectral characterization

Photophysical characterization was performed as previously described^33,57^. Absorption spectra were recorded with a Lambda900 spectrophotometer (PerkinElmer), and fluorescence spectra with an LS55 spectrofluorimeter (PerkinElmer). Extinction coefficients and fractional concentrations of anionic and neutral forms (free and bound) were determined by alkaline titration. Fluorescence quantum yields were measured using an integrating sphere (Quantaurus-QY, Hamamatsu) with 485–505 nm excitation. Fluorescence lifetimes were obtained with a Digital Frequency Domain ChronosDFD system (ISS) using 515-nm excitation/520LP emission (Zn^2+^-saturated) and 445-nm excitation/460LP or 525/40 emission (Zn^2+^-free). Two-photon excitation spectra and cross sections were recorded with a DeepSee Insight laser (MKS-SpectraPhysics) and PC1 photon-counting spectrofluorimeter (ISS). Excitation spectra employed 770SP, 633SP, and 535/50 emission filters; cross-section measurements used 960-nm two-photon and 488-nm one-photon excitation with Rhodamine 6G as a standard, collecting fluorescence through 770SP and 520LP filters.

### Mammalian cell culture, transfection, and characterization

Gene fragments were PCR-amplified and cloned into the pDisplay vector between Bgl II and Sal I sites. HEK293T cells were maintained in DMEM (4.5 g/L glucose, 10% FBS), and SH-SY5Y cells in DMEM/F12 (10% FBS), at 37 °C with 5% CO_2_. Cells in 35-mm dishes were transfected as previously described^33^. HEK293T cells received 1 μg plasmid DNA and 3 μg PEI (linear, 25 kDa; Polysciences), and SH-SY5Y cells received 1 μg DNA with 3 μg Lipofectamine 2000 (Thermo Fischer). Before imaging, cells were rinsed with DPBS and equilibrated for 30 min in Mammalian Cell Imaging Buffer (138 mM NaCl, 2.2 mM KCl, 2 mM glucose, 2 mM CaCl_2_, 2 mM MgCl_2_, 25 mM HEPES, pH 7.4). Images were acquired on a Leica DMi8 microscope with a 40× oil objective. A confocal module was used to analyze localization, and for co-expression experiments, sequential imaging used 488-nm (500–550 nm emission) and 532-nm (580–700 nm emission) channels. Single-color wide-field time-lapse imaging used a FITC filter set (480/40 nm excitation, 535/50 nm emission). Fluorescence lifetime imaging was performed with a Lambert LIFA-Toggle frequency-domain FLIM system using a 474-nm LED, 480/40 nm excitation and 527/30 nm emission filters, 1 s exposure, 40 MHz modulation, and 12 phase steps. Fluorescein (τ = 4.1 ns) served as the lifetime reference. The fluorescence lifetime values reported in this work were derived from phase shifts.

### Viral preparation and titration

AAV was produced as previously described^33^. Briefly, HEK293T cells were co-transfected with pAAV-hSyn-hpGRISZ-TM, pAdDeltaF6, and pAAV2/9n, and maintained in DMEM (4.5 g/L glucose, 4% FBS) supplemented with 0.1 M sucrose and 10 mM HEPES for 96 hours. Cells were collected by centrifugation, and viral particles were isolated from both the culture supernatant and cell pellet lysates. Viral purification was performed by gradient ultracentrifugation, followed by buffer exchange into DPBS containing 0.001% pluronic and 200 mM NaCl. The purified virus was aliquoted, flash-frozen, and stored at −80 °C. Viral titers were quantified by qPCR.

### Wide-field imaging of auditory cortex and data analysis

Surgery and wide-field imaging were performed largely as previously described^33^, with modifications to assess hpGRISZ-TM responses. Mice were injected with AAVs (500 nL, 5.5 × 10^13^ gc/mL) into the right ACtx at P42–49 and imaged at P63–70. Twelve to twenty-five days post-injection, a ~1 mm craniotomy was made over A1 under anesthesia. A glass micropipette containing ACSF with 100 μM ZX1 was inserted at the craniotomy edge using a micromanipulator, and mice were allowed to recover from isoflurane for 60 min before imaging. Broadband acoustic stimuli (6–64 kHz, 100 ms, 60–70 dB SPL) were delivered in a calibrated sound-attenuating chamber, with sound delivery and image acquisition synchronized using Ephus (MATLAB). Wide-field hpGRISZ-TM fluorescence was recorded with epifluorescence optics (EGFP filter set), a 4× objective, and a CCD camera at 20 Hz with 4× spatial binning (174 × 130 pixels; 171.1 μm^2^ per pixel). Each stimulus was repeated 8–10 times. After the initial recordings, ZX1 (100 μM) or ACSF was infused at 30 nL/min for 20 min, then at 9 nL/min, and responses were re-measured. For analysis, normalized fluorescence changes were calculated at each pixel as ΔF/F = (F – F_0_)/F_0_, where F_0_ was the mean fluorescence during 2–3 s before stimulus onset. To identify sound-responsive regions, ΔF/F movies were filtered with a 4th-order 5 Hz low-pass Butterworth filter, then temporally averaged across 500 ms (10 consecutive frames) after stimulus onset. Quantification of sound-evoked responses was performed by centering a 30 × 30 pixel ROI on the region with the largest ΔF/F amplitude; traces were calculated as mean fluorescence intensity within the ROI normalized to F_0_. To spatially segregate signal excursion components, ΔF/F was also computed within 10 × 10 pixel ROIs swept across the field at 5-pixel increments, and traces were clustered by pairwise Euclidean distance to capture temporal similarity. To evaluate potential motion artifacts, image series were normalized to 8-bit, smoothed with a Gaussian blur (5 × 5 kernel), and processed by Canny edge detection (OpenCV). A blood vessel-containing ROI was selected, and average displacement of edge pixels (Y-coordinate change relative to two frames earlier, 100 ms) was quantified across frames.

### In vivo two-photon imaging of auditory cortex and data analysis

Two-photon imaging of L2/3 A1 neurons (~200 μm below pia) in awake, head-fixed mice was performed as previously described^17,44^, using a galvanometer-based microscope (Scientifica) with a 40×, 0.8 NA objective (Olympus) and a mode-locked Ti:Sapphire laser (940 nm, MaiTai HP, Newport). Green fluorescence was collected with a PMT and a Semrock FF03-525/50 filter. Images were acquired at 5 Hz over a 145 × 145 μm field (256 × 256 pixels). Broadband sounds (6–64 kHz, 100 ms, 20–80 dB SPL in 10 dB steps, pseudo-random order) were delivered every 7 s, and imaging sessions lasted 20–30 min. Images were analyzed post hoc using a custom program, and open-source routines, written using Python 3.5 and MATLAB as described previously^17,43,44,58^. Motion correction was applied using NoRMCorre^59^, and somatic regions of interest (ROIs) were selected from temporal averages of all image frames. Fluorescence signals were extracted, and neuropil contamination was corrected using FISSA^60^, with the neuropil vector weighted by 0.8 as described previously^43,58^. Corrected somatic fluorescence values (F) were converted to ΔF/F using the baseline fluorescence during the 1 s pre-stimulus window. For each neuron, ΔF/F responses were averaged across 8–10 trials per sound. Responses were considered significant if post-stimulus ΔF/F exceeded baseline mean + 3 SD. Peak amplitudes were measured within 300–1300 ms after stimulus onset. Response threshold was defined as the lowest of at two consecutive levels which elicited a significant response; only neurons with thresholds ≤70 dB SPL were included in further analysis.

### Surgeries, fiber photometry, and analysis of BLA Zn^2+^ dynamics

C57BL/6J mice (2–5 months) were injected with 400 nL AAV (5.5 × 10^13^ gc/mL) into the right BLA (AP: −1.6 mm; ML: +3.3 mm; DV: −4.75 mm) at a rate of 1 nL/s under sevoflurane anesthesia. A hybrid cannula (Cat # FI_OmFC-ZF_140/190_6.0mm, Doric Lenses) was implanted 0.1 mm above the target for combined pharmacology and photometry. Mice received perioperative buprenorphine and recovered ≥4 weeks before experiments. Zn^2+^ dynamics in the BLA were recorded with a dual-wavelength fiber photometry system (Doric Lenses). Excitation light (465 nm, Zn^2+^ channel; 405 nm, isosbestic control) was routed via a minicube to the implanted fiber optic cannula (400 μm core, 0.50 NA). Fluorescence was collected through the same fiber, separated by emission filters, and detected with a Newport 2151 photoreceiver.

Signals were acquired with Doric Studio Software, synchronized to behavioral events via TTL inputs. For pharmacological manipulations, 50 mM of ZX1 (Cat # 07-0350, Strem Chemicals) stock solution was diluted to 200 μM in PBS. ZX1 or normal saline (2 μL) was infused intracranially through the hybrid cannula immediately before behavioral tests using a 5-μL Hamilton syringe. Behavioral assays were conducted in custom-built operant chambers (17.8 × 19 × 23 cm) controlled by pyControl. Three paradigms were used: (1) Looming sound^48^— 20 trials of a rising-intensity stimulus (20–70 dB over 0.4 s, repeated 10 times/trial, ITI 60–75 s) after 20 s habituation; (2) Pure tone (20 kHz)^49^— 20 trials of a 20 kHz, 100 dB tone (3 s, ITI 30– 40 s) after habituation; (3) Foot shock— 10 trials of mild shock (0.25 mA, 1 s, ITI 20–35 s) after habituation. Fiber photometry data were analyzed using GuPPy (RRID:SCR_022353). Signals were motion-corrected, normalized to the isosbestic channel, and converted to ΔF/F. Event-aligned post-stimulus time histograms (PSTHs) were generated (looming: −10 to +30 s; pure tone: −5 to +15 s; shock: −5 to +25 s). Fluorescence values were z-scored to baseline, and stimulus-evoked activity quantified as the area under the curve (AUC). Pre- and stimulus intervals were defined as follows: looming (−3–0 s vs. 0–3 s), pure tone (−2–0 s vs. 0–2 s), and foot shock (−2–0 s vs. 0–6 s).

### Statistical analysis

Statistical analyses were performed in GraphPad Prism. For fiber photometry experiments, outliers were identified using box-and-whisker plots with Tukey’s test and removed before further analysis. Data normality was assessed with the Shapiro–Wilk test. For all experiments, results are reported as mean ± SD or mean ± SEM, with the significance threshold set at p ≤ 0.05. Statistical details, sample sizes, and test types are provided in the figure legends.

### Data Availability

The plasmids pDisplay-hpGRISZ and pAAV-hSyn-hpGRISZ-TM and their sequence information will be deposited to Addgene. All key data and experimental methods are presented in the main text or the supplementary materials. Source data will be provided with this paper.

## Acknowledgements

This work was supported by the National Institutes of Health (R01EB033172, H.A. and T.T.; R01EB035430, H.A.; R01AG077773, H.A.; R01-DC019618, T.T; R01-DC020923, T.T.; U24NS109107, M.D.); the Hearing Health Foundation (1307587, M.K.); the European Research Council (101003187; A.J.R.); the “la Caixa” Foundation (LCF/PR/HR20/52400020; A.J.R.); the Portuguese Foundation for Science and Technology (FCT) (PTDC/MED-NEU/4804/2020, A.J.R.; CEECIND/CP2841/CT0023, C.S.C.; 2022.12973.BD, R.C.); and the Bial Foundation and the Órdem dos Médicos (the Maria de Sousa Award, C.S.-C.).

## Author contributions

H.A. provided overall project administration. H.Z. engineered the hpGRISZ sensor and performed most in vitro and cellular characterization, with assistance from W.H., Y.Z., and S.Z. H.Z., W.H., and X.T. prepared the virus. T.T. directed experimental design and management for M.K., S.Y., and P.A.C. in wide-field and 2P imaging of the ACtx. A.J.R. directed experimental design and management for R.C. and C.S.-C. in fiber photometry recording of the BLA. M.D. performed photophysical characterization using proteins prepared by H.Z. Data analysis and curation were carried out by H.Z., M.K., S.Y., P.A.C., R.C., C.S.-C., and M.D. H.A., H.Z., M.K., P.A.C., C.S.-C., M.D., T.T., and A.J.R. wrote the manuscript with input from all authors.

Funding was acquired by H.A., T.T., A.J.R. and M.D. for research conducted in their own laboratories.

## Competing interests

The authors declare that they have no competing interests.

## Notes

### Competing Interest Statement

The authors have declared no competing interest.

## References

1. Andreini, C., Banci, L., Bertini, I. & Rosato, A. Counting the zinc-proteins encoded in the human genome. J Proteome Res 5, 196–201 (2006).

2. Lo, M.N., Damon, L.J., Wei Tay, J., Jia, S. & Palmer, A.E. Single cell analysis reveals multiple requirements for zinc in the mammalian cell cycle. Elife 9, e51107 (2020).

3. Krall, R.F., Tzounopoulos, T. & Aizenman, E. The function and regulation of zinc in the brain. Neuroscience 457, 235–258 (2021).

4. Wessels, I., Maywald, M. & Rink, L. Zinc as a Gatekeeper of Immune Function. Nutrients 9 (2017).

5. Ahmad, R., Shaju, R., Atfi, A. & Razzaque, M.S. Zinc and Diabetes: A Connection between Micronutrient and Metabolism. Cells 13 (2024).

6. Li, Y.V. Zinc and insulin in pancreatic beta-cells. Endocrine 45, 178–189 (2014).

7. Li, Z., Liu, Y., Wei, R., Yong, V.W. & Xue, M. The important role of zinc in neurological diseases. Biomolecules 13, 28 (2022).

8. Lee, E.-J. et al. Trans-synaptic zinc mobilization improves social interaction in two mouse models of autism through NMDAR activation. Nat Commun 6, 7168 (2015).

9. Swardfager, W. et al. Zinc in depression: a meta-analysis. Biol Psychiatry 74, 872–878 (2013).

10. Doboszewska, U. et al. Zinc signaling and epilepsy. Pharmacol Ther 193, 156–177 (2019).

11. Adlard, P.A. & Bush, A.I. Metals and Alzheimer’s disease. J Alzheimers Dis 10, 145–163 (2006).

12. Pan, E. et al. Vesicular zinc promotes presynaptic and inhibits postsynaptic long-term potentiation of mossy fiber-CA3 synapse. Neuron 71, 1116–1126 (2011).

13. Upmanyu, N. et al. Colocalization of different neurotransmitter transporters on synaptic vesicles is sparse except for VGLUT1 and ZnT3. Neuron 110, 1483–1497.e1487 (2022).

14. Bender, P.T., McCollum, M., Boyd-Pratt, H., Mendelson, B.Z. & Anderson, C.T. Synaptic zinc potentiates AMPA receptor function in mouse auditory cortex. Cell Rep 42 (2023).

15. Brown, C.E. & Dyck, R.H. Modulation of synaptic zinc in barrel cortex by whisker stimulation. Neuroscience 134, 355–359 (2005).

16. Krall, R.F. et al. Synaptic zinc inhibition of NMDA receptors depends on the association of GluN2A with the zinc transporter ZnT1. Science Adv 6, eabb1515 (2020).

17. Anderson, C.T., Kumar, M., Xiong, S. & Tzounopoulos, T. Cell-specific gain modulation by synaptically released zinc in cortical circuits of audition. Elife 6 (2017).

18. Kouvaros, S., Kumar, M. & Tzounopoulos, T. Synaptic Zinc Enhances Inhibition Mediated by Somatostatin, but not Parvalbumin, Cells in Mouse Auditory Cortex. Cereb Cortex 30, 3895–3909 (2020).

19. Pratt, E.P.S., Damon, L.J., Anson, K.J. & Palmer, A.E. Tools and techniques for illuminating the cell biology of zinc. Biochim Biophys Acta Mol Cell Res 1868, 118865 (2021).

20. Fang, H. et al. Recent advances in Zn2+ imaging: From organelles to in vivo applications. Curr Opin Chem Biol 76, 102378 (2023).

21. Nolan, E.M. & Lippard, S.J. Small-molecule fluorescent sensors for investigating zinc metalloneurochemistry. Acc Chem Res 42, 193–203 (2009).

22. Li, D., Liu, L. & Li, W.H. Genetic targeting of a small fluorescent zinc indicator to cell surface for monitoring zinc secretion. ACS Chem Biol 10, 1054–1063 (2015).

23. Gest, A.M.M. et al. Molecular Spies in Action: Genetically Encoded Fluorescent Biosensors Light up Cellular Signals. Chem Rev 124, 12573–12660 (2024).

24. Carter, K.P., Young, A.M. & Palmer, A.E. Fluorescent sensors for measuring metal ions in living systems. Chem Rev 114, 4564–4601 (2014).

25. Park, J.G., Qin, Y., Galati, D.F. & Palmer, A.E. New sensors for quantitative measurement of mitochondrial Zn2+. ACS Chem Biol 7, 1636–1640 (2012).

26. Hessels, A.M. et al. eZinCh-2: a versatile, genetically encoded FRET sensor for cytosolic and intraorganelle Zn2+ imaging. ACS Chem Biol 10, 2126–2134 (2015).

27. Vinkenborg, J.L. et al. Genetically encoded FRET sensors to monitor intracellular Zn2+ homeostasis. Nat Methods 6, 737–740 (2009).

28. Hessels, A.M. & Merkx, M. Genetically-encoded FRET-based sensors for monitoring Zn2+ in living cells. Metallomics 7, 258–266 (2015).

29. Minckley, T.F. et al. Sub-nanomolar sensitive GZnP3 reveals TRPML1-mediated neuronal Zn2+ signals. Nat Commun 10, 4806 (2019).

30. Chen, M. et al. Genetically encoded, photostable indicators to image dynamic Zn2+ secretion of pancreatic islets. Anal Chem 91, 12212–12219 (2019).

31. Dischler, A.M., Maslar, D., Zhang, C. & Qin, Y. Development and Characterization of a Red Fluorescent Protein-Based Sensor RZnP1 for the Detection of Cytosolic Zn2+. ACS Sensors 7, 3838–3845 (2022).

32. Qin, Y., Sammond, D.W., Braselmann, E., Carpenter, M.C. & Palmer, A.E. Development of an optical Zn2+ probe based on a single fluorescent protein. ACS Chem Biol 11, 2744–2751 (2016).

33. Wu, T. et al. A genetically encoded far-red fluorescent indicator for imaging synaptically released Zn2+. Sci Adv 9, eadd2058 (2023).

34. Chen, Z. & Ai, H.W. Single Fluorescent Protein-Based Indicators for Zinc Ion (Zn2+). Anal Chem 88, 9029–9036 (2016).

35. Kodirov, S.A. et al. Synaptically released zinc gates long-term potentiation in fear conditioning pathways. Proc Natl Acad Sci U S A 103, 15218–15223 (2006).

36. McAllister, B.B. & Dyck, R.H. Zinc transporter 3 (ZnT3) and vesicular zinc in central nervous system function. Neurosci Biobehav Rev 80, 329–350 (2017).

37. Hopfner, K.P. et al. The Rad50 zinc-hook is a structure joining Mre11 complexes in DNA recombination and repair. Nature 418, 562–566 (2002).

38. Chen, Z., Zhang, S., Li, X. & Ai, H.W. A high-performance genetically encoded fluorescent biosensor for imaging physiological peroxynitrite. Cell Chem Biol 28, 1542–1553.e1545 (2021).

39. Pédelacq, J.D., Cabantous, S., Tran, T., Terwilliger, T.C. & Waldo, G.S. Engineering and characterization of a superfolder green fluorescent protein. Nat Biotechnol 24, 79–88 (2006).

40. Linton, E., Walsh, M.K., Sims, R.C. & Miller, C.D. Translocation of green fluorescent protein by comparative analysis with multiple signal peptides. Biotechnol J 7, 667–676 (2012).

41. Zhao, Y. et al. An expanded palette of genetically encoded Ca²? indicators. Science 333, 1888–1891 (2011).

42. Kumar, M., Xiong, S., Tzounopoulos, T. & Anderson, C.T. Fine Control of Sound Frequency Tuning and Frequency Discrimination Acuity by Synaptic Zinc Signaling in Mouse Auditory Cortex. J Neurosci 39, 854–865 (2019).

43. Cody, P.A. & Tzounopoulos, T. Neuromodulatory mechanisms underlying contrast gain control in mouse auditory cortex. J Neurosci, JN-RM-2054-2021 (2022).

44. Cody, P., Kumar, M. & Tzounopoulos, T. Cortical Zinc Signaling Is Necessary for Changes in Mouse Pupil Diameter That Are Evoked by Background Sounds with Different Contrasts. J Neurosci 44 (2024).

45. McCollum, M., Manning, A., Bender, P.T.R., Mendelson, B.Z. & Anderson, C.T. Cell-type-specific enhancement of deviance detection by synaptic zinc in the mouse auditory cortex. Proc Natl Acad Sci U S A 121, e2405615121 (2024).

46. Martel, G., Hevi, C., Friebely, O., Baybutt, T. & Shumyatsky, G.P. Zinc transporter 3 is involved in learned fear and extinction, but not in innate fear. Learn Mem 17, 582–590 (2010).

47. Cole, T.B., Martyanova, A. & Palmiter, R.D. Removing zinc from synaptic vesicles does not impair spatial learning, memory, or sensorimotor functions in the mouse. Brain Res 891, 253–265 (2001).

48. Li, Z. et al. Corticostriatal control of defense behavior in mice induced by auditory looming cues. Nat Commun 12, 1040 (2021).

49. Moriya, S. et al. Acute Aversive Stimuli Rapidly Increase the Activity of Ventral Tegmental Area Dopamine Neurons in Awake Mice. Neuroscience 386, 16–23 (2018).

50. Blakemore, L.J. & Trombley, P.Q. Zinc as a Neuromodulator in the Central Nervous System with a Focus on the Olfactory Bulb. Front Cell Neurosci 11, 297 (2017).

51. Izumi, Y., Auberson, Y.P. & Zorumski, C.F. Zinc modulates bidirectional hippocampal plasticity by effects on NMDA receptors. J Neurosci 26, 7181–7188 (2006).

52. Barberis, A., Cherubini, E. & Mozrzymas, J.W. Zinc inhibits miniature GABAergic currents by allosteric modulation of GABAA receptor gating. J Neurosci 20, 8618–8627 (2000).

53. Kouvaros, S. et al. A CRE/DRE dual recombinase transgenic mouse reveals synaptic zinc-mediated thalamocortical neuromodulation. Sci Adv 9, eadf3525 (2023).

54. Maret, W. Analyzing free zinc(II) ion concentrations in cell biology with fluorescent chelating molecules. Metallomics 7, 202–211 (2015).

55. Bizup, B. & Tzounopoulos, T. On the genesis and unique functions of zinc neuromodulation. J Neurophysiol 132, 1241–1254 (2024).

56. Shindo, Y. et al. Genetically encoded sensors for analysing neurotransmission among synaptically-connected neurons. bioRxiv, 2022.2004.2003.486903 (2022).

57. Drobizhev, M., Molina, R.S. & Hughes, T.E. Characterizing the Two-photon Absorption Properties of Fluorescent Molecules in the 680-1300 nm Spectral Range. Bio Protoc 10 (2020).

58. Kerlin, A.M., Andermann, M.L., Berezovskii, V.K. & Reid, R.C. Broadly tuned response properties of diverse inhibitory neuron subtypes in mouse visual cortex. Neuron 67, 858– 871 (2010).

59. Pnevmatikakis, E.A. & Giovannucci, A. NoRMCorre: An online algorithm for piecewise rigid motion correction of calcium imaging data. J Neurosci Methods 291, 83–94 (2017).

60. Keemink, S.W. et al. FISSA: A neuropil decontamination toolbox for calcium imaging signals. Sci Rep 8, 3493 (2018).

