## Supplementary Information for "hpGRISZ: a high-performance fluorescent biosensor for in vivo imaging of synaptic zinc dynamics"

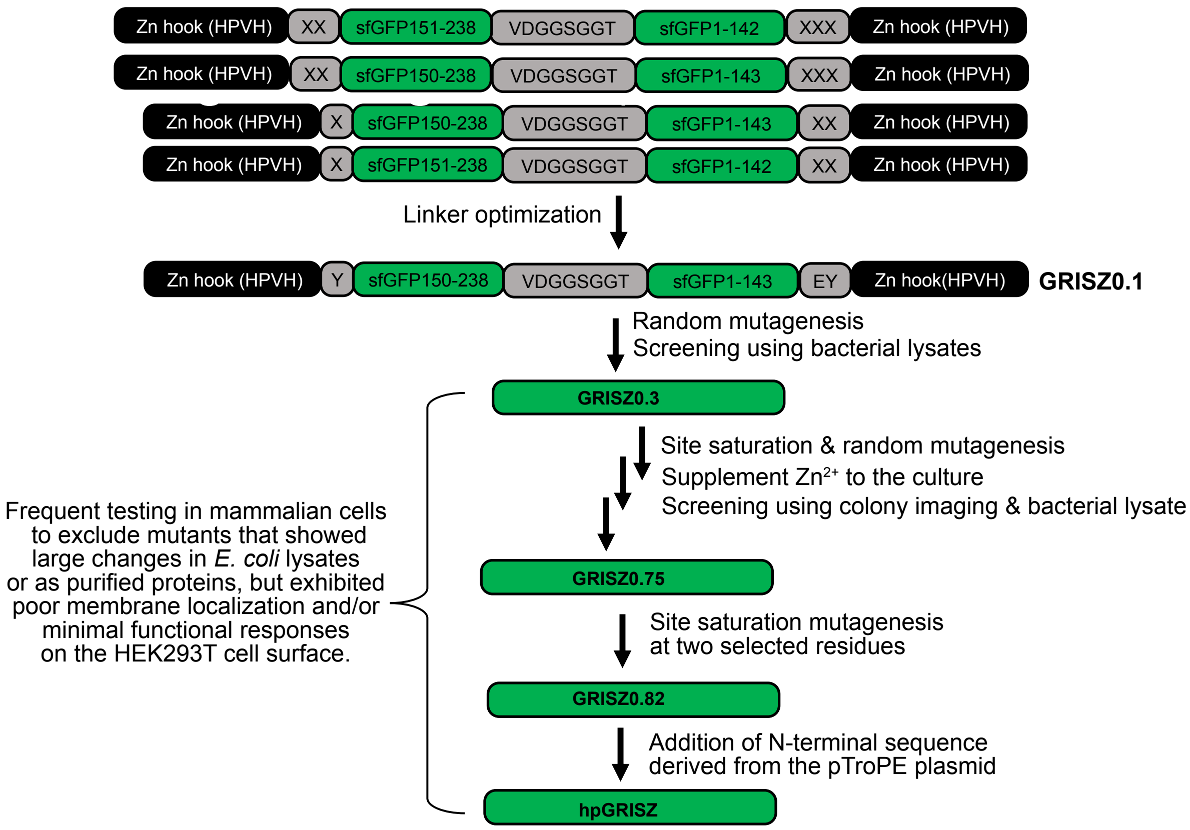


**Figure S1. Illustration of the multi-step process used to engineer hpGRISZ.**

Schematic flowchart illustrating the design and optimization of hpGRISZ. Modified zinc hook (HPVH) motifs were fused to the N- and C-termini of circularly permuted sfGFP via variable linker sequences (X = 20 amino acids encoded using the NNK codon). Linker optimization produced the initial variant GRISZ0.1, followed by iterative mutagenesis and library screening to generate progressively improved intermediates (up to GRISZ0.75). Subsequently, screening of a small library with saturation mutagenesis at two selected residues yielded GRISZ0.82. Incorporation of the N-terminal sequence from the pTroPE plasmid proved essential for high responses at the mammalian cell surface, resulting in the final construct, hpGRISZ.

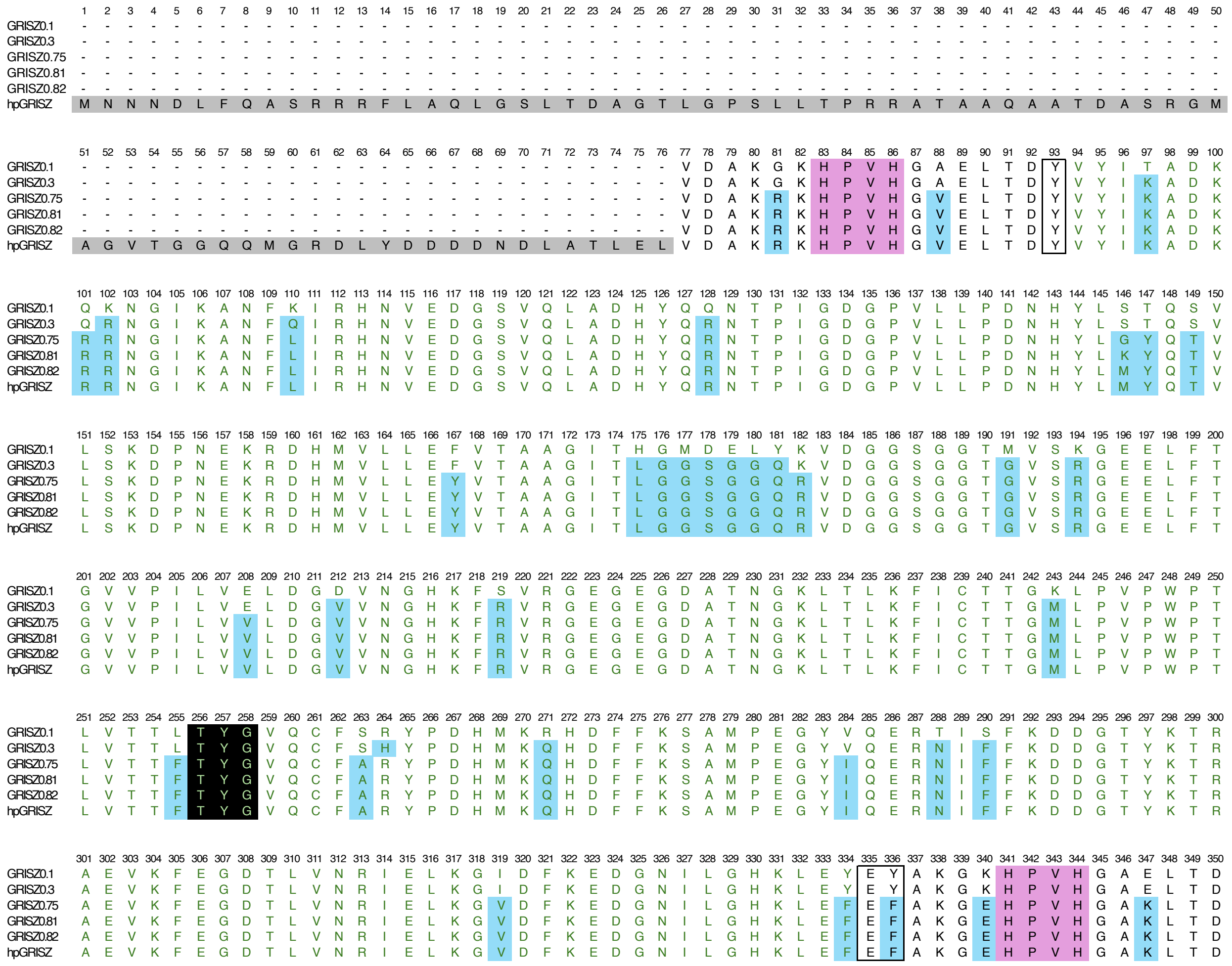

**Figure S2. Sequence alignment of hpGRISZ with earlier variants.**

Sequences derived from sfGFP (residues 94–334) are shown in green. The chromophore-forming residues (256–258) are shaded black. Zinc hook motifs (residues 83–86 and 341–344) are shaded magenta. Linkers connecting cpsfGFP and the zinc hooks (residues 93 and 335/336) are outlined in black boxes. Mutations acquired during indicator engineering after GRISZ0.1 are shaded cyan.

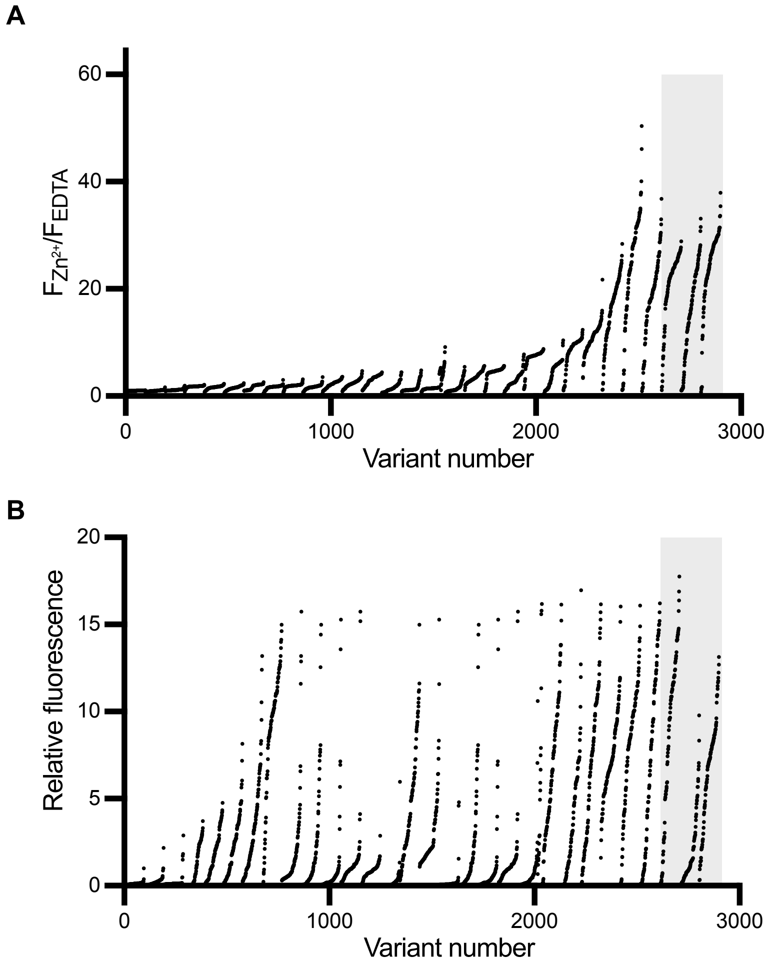


**Figure S3. Fluorescence properties of GRISZ variants characterized in plate reader–based cell lysate assays.**

**(A)** Fluorescence response to $Zn^{2+}$ for characterized variants, calculated as the fluorescence with $Zn^{2+}$ divided by the fluorescence with EDTA. **(B)** Relative fluorescence intensities of the variants in the $Zn^{2+}$-bound state. Screening was performed with 250 µM $ZnCl_2$, except for the final three rounds (variants 2613–2900, shaded in grey), where 50 µM $ZnCl_2$ was used. The graphs include all variants selected for plate reader-based cell lysate assays before the development of GRISZ0.82, which was obtained through saturation mutagenesis of two sites in GRISZ0.75.

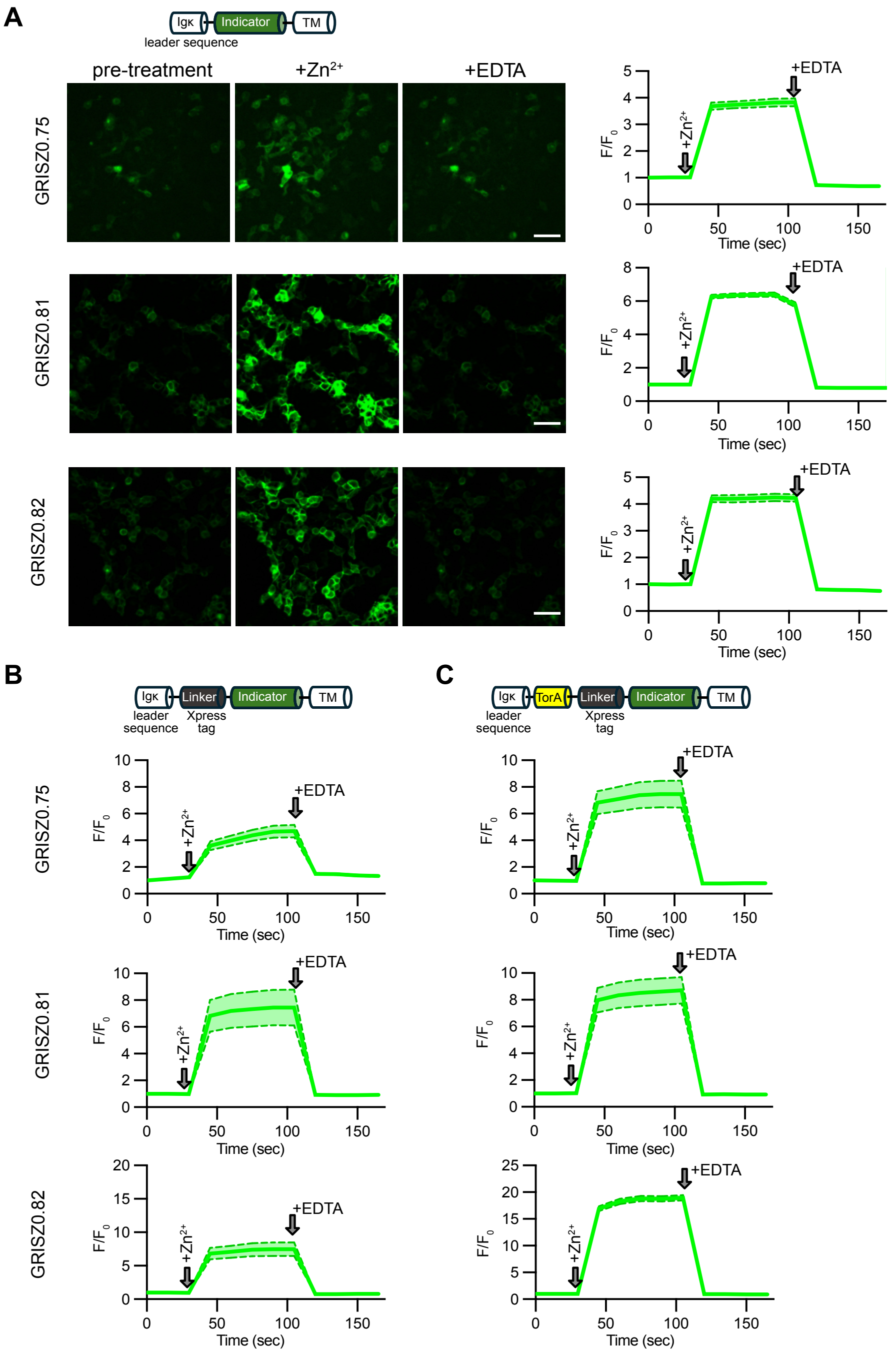


**Figure S4. Characterization of GRISZ construct variants (GRISZ0.75, GRISZ0.81, and GRISZ0.82) expressed at the surface of HEK293T cells.**

(**A**) Representative fluorescence images and response traces of GRISZ variants incorporated into pDisplay. Scale bar, 25 μm. Data are presented as mean ± SEM (n = 23, 54, and 51 cells, respectively). (**B**) Fluorescence response traces when a linker (an Xpress Tag derived from pTorPE) was appended to the N-termini of the sensors in pDisplay. Data are presented as mean ± SEM (n = 30, 7, and 10 cells, respectively). (**C**) Fluorescence response traces when the TorA and linker sequence (derived from pTorPE) was appended to the N-termini of the sensors in pDisplay. Data are presented as mean ± SEM (n = 10, 11, and 36 cells, respectively).

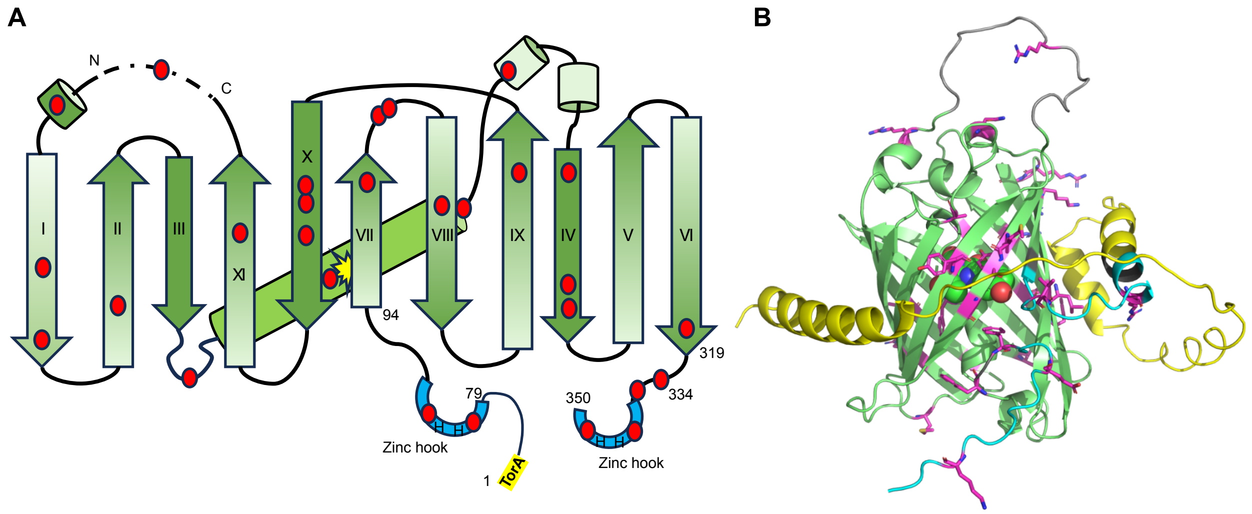


**Figure S5. Locations of mutations acquired during the sensor engineering process (from GRISZ0.1 to hpGRISZ).**

(**A**) Mutations mapped onto the secondary structure of hpGRISZ. Arrows represent β-strands, cylinders represent α-helices, and cyan semicircles indicate the Rad50-derived cysteine-less zinc hooks (with "H" denoting histidine). Point mutations acquired during the engineering process are shown as red ovals. (**B**) Mutations mapped onto the AlphaFold3-predicted 3D structure of hpGRISZ ($Zn^{2+}$-free state). AlphaFold3 did not yield a reasonable model of $Zn^{2+}$-bound hpGRISZ. Structural features are highlighted as follows: yellow, TorA and Xpress sequences; cyan, cysteine-less zinc hook; green, circularly permuted superfolder GFP; gray, linker connecting the original GFP N- and C-termini; magenta, mutations acquired during the engineering process.

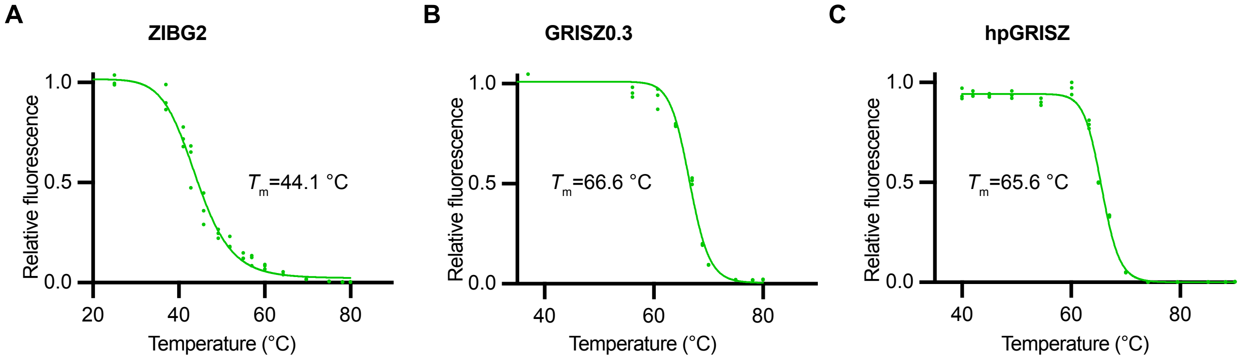


**Figure S6. Thermostability characterization of the indicated GEZI variants.**

Purified proteins (5 μM) in HEPES buffer containing 100 μM EDTA were incubated at the indicated temperatures for 1 h in a thermal cycler. Fluorescence intensities were measured immediately afterward using a plate reader.

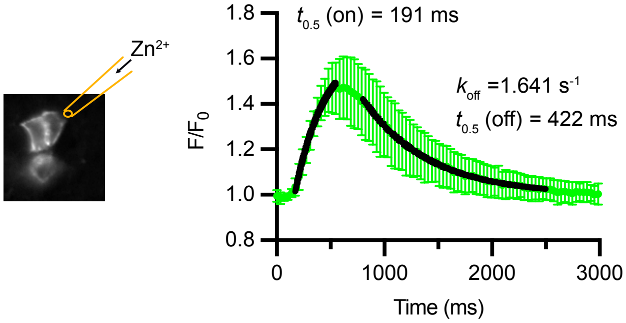


**Figure S7. Response kinetics characterization of hpGRISZ-TM.**

Response of hpGRISZ expressed on the surface of HEK293T cells to puff application of $Zn^{2+}$. Data are presented as mean ± SD from 15 trials (three cells, five applications each) and fitted with monoexponential functions for both onset and decay. The derived on- and off-half-times, as well as the $k_{off}$ values, fall within the same range as those previously reported for FRISZ.

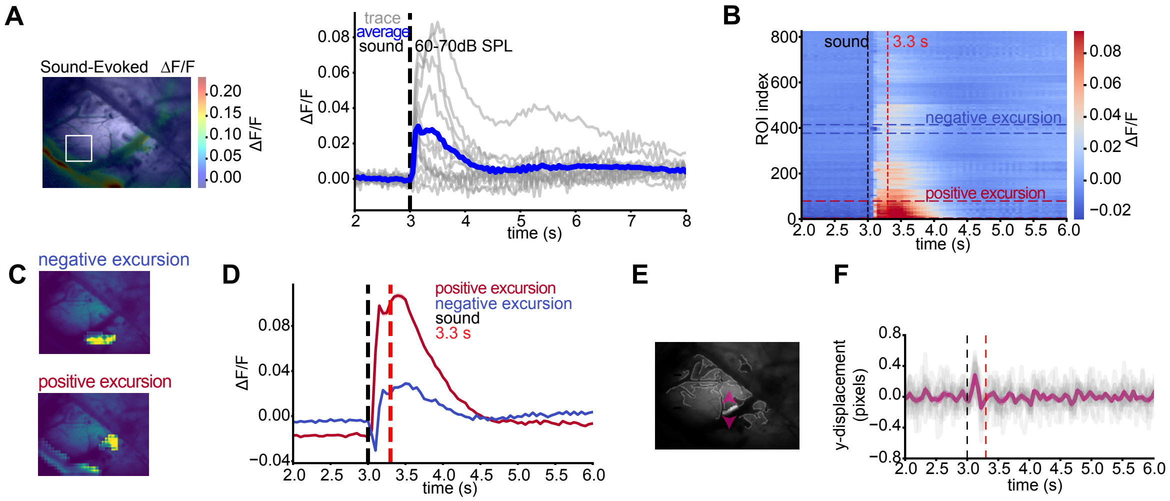


**Figure S8. Wide-field imaging of sound-evoked $Zn^{2+}$ responses in auditory cortex and correction of motion artifacts.**

**(A)** Representative wide-field ΔF/F response to a 100 ms broadband sound (6–64 kHz, 60–70 dB). *Left:* grayscale image of the ACtx through the craniotomy with ROI indicated (white square). *Right:* ΔF/F traces from the ROI (gray, individual; blue, average), aligned to sound onset at 3 s (dotted line). **(B)** ΔF/F traces from 10×10-pixel ROIs, sorted by temporal similarity. A prominent negative signal excursion (blue dotted box) ends by 3.3 seconds (red dotted line), while a prominent positive excursion is outlined in ruby dotted lines. **(C)** Spatial maps of ΔF/F averaged from negative (blue) or positive (ruby) ROIs identified in panel B; non-ROI regions darkened. **(D)** Average ΔF/F traces corresponding to negative (blue) and positive (ruby) signal excursions. **(E)** Edge detection of the imaging field in panel A. Magenta arrows enclose lightened ROI of tracked pixels along blood vessel edge. **(F)** Y-axis displacement of the ROI edge shown in panel E, calculated as average difference in y-axis coordinate of edge labeled pixels from 2 frames prior. Individual (gray) and average (magenta) traces are shown.

### Table S1 | Summary of photophysical properties of hpGRISZ.

| | hpGRISZ | |
|---|---|---|
| | $Zn^{2+}$-free | $Zn^{2+}$-bound |
| Peak Excitation $\lambda_{ex}$ (nm) | 501 | 499 |
| Peak Emission $\lambda_{em}$ (nm) | 520 | 516 |
| $F/F_0$ | 25 | |
| $K_d$ (μM) [a] | 12 | |
| Apparent Hill coefficient ($n_H$) | 1.6 | |
| Fluorescence lifetime $\tau$ (ns) [b] | 2.92 (83%)<br>0.56 (17%) | 3.37 |
| Effective extinction coefficient ε ($mM^{-1}$ $cm^{-1}$) [c] | 1.48 | 25.6 |
| Quantum yield φ | 0.66 | 0.92 |
| 1P brightness (ε×φ) | 0.97 | 23.6 |
| 2P brightness (GM) at the indicated excitation wavelength | 0.35 (940 nm) | 10 (940 nm) |
| $F/F_0$ at indicated 2P excitation wavelength | 29 (940 nm)<br>33 (996 nm) | |

[a] Determined using purified protein and intensity measurements. When displayed on the HEK293T cell surface, the sensor showed a $K_d$ of 4.4 μM ($n_H$= 1.6) from intensity measurements and 0.82 μM ($n_H$= 1.1) from frequency-domain fluorescence lifetime measurements.

[b] Determined using purified protein and time-domain measurements. When displayed on the HEK293T cell surface, the sensor exhibited a lifetime change from 3.79 ns ($Zn^{2+}$-free) to 4.28 ns ($Zn^{2+}$-bound) as measured by frequency-domain fluorescence lifetime analysis.

[c] Presented as per mM of the total protein (a product of the relative fraction and the true extinction coefficient for per mM of the anionic chromophore).
